# Generalization of Cortical MOSTest Genome-Wide Associations Within and Across Samples

**DOI:** 10.1101/2021.04.23.441215

**Authors:** Robert J. Loughnan, Alexey A. Shadrin, Oleksandr Frei, Dennis van der Meer, Weiqi Zhao, Clare E. Palmer, Wesley K. Thompson, Carolina Makowski, Terry L. Jernigan, Ole A. Andreassen, Chun Chieh Fan, Anders M. Dale

## Abstract

Genome-Wide Association studies have typically been limited to single phenotypes, given that high dimensional phenotypes incur a large multiple comparisons burden: ~1 million tests across the genome times the number of phenotypes. Recent work demonstrates that a Multivariate Omnibus Statistic Test (MOSTest) is well powered to discover genomic effects distributed across multiple phenotypes. Applied to cortical brain MRI morphology measures, MOSTest has resulted in a drastic improvement in power to discover loci when compared to established approaches (min-P). One question that arises is how well these discovered loci replicate in independent data. Here we perform 10 times cross validation within 35,644 individuals from UK Biobank for imaging measures of cortical area, thickness and sulcal depth (>1,000 dimensionality for each). By deploying a replication method that aggregates discovered effects distributed across multiple phenotypes, termed PolyVertex Score (PVS), we demonstrate a higher replication yield and comparable replication rate of discovered loci for MOSTest (# replicated loci: 348-845, replication rate: 94-95%) in independent data when compared with the established min-P approach (# replicated loci: 31-68, replication rate: 65-80%). An out-of-sample replication of discovered loci was conducted with a sample of 8,336 individuals from the Adolescent Brain Cognitive Development^®^ (ABCD) study, who are on average 50 years younger than UK Biobank individuals. We observe a higher replication yield and comparable replication rate of MOSTest compared to min-P. This finding underscores the importance of using well-powered multivariate techniques for both discovery and replication of high dimensional phenotypes in Genome-Wide Association studies.

## Introduction

Performing Genome Wide Association Studies (GWAS) on high dimensional phenotypes incurs a large multiple comparisons burden (number of independent genetic tests by number of phenotypes) using traditional approaches, which can result in low power to detect associations. Vertex-wise measures of cortical morphology (area, thickness and sulcal depth) represent high dimensional phenotypes (>1000 dimensions) and, from twin studies, are known to have high heritabilities of up to 90% and 50% for total and regional area respectively, and 80% and 60% for mean and regional thickness respectively(1,2). Our group has previously developed a novel Multivariate Omnibus Test (MOSTest) (3–5), which aggregates the effect of a genomic variant across the cortex. This method significantly boosts discovery of genetic loci linked to cortical morphology, with an up to 10x increase in number of loci discovered – when compared to an established approach (min-P) deployed for the same phenotypes(5). Additionally, discovered loci show strong enrichment with pathways involved in neurogenesis and cell differentiation. Two main benefits of MOSTest over established techniques, like min-P, are: 1) its ability to aggregate pleiotropic effects into a single statistical test and 2) it drastically reduces the multiple comparison burden across the dimensionality of phenotypes, while still accounting for genome-wide multiple comparisons correction. Given such a dramatic increase in discovery of genomic loci, it is of interest to understand how well these discoveries replicate in independent data.

Here we perform 10-times cross validation with brain imaging data taken from the UK Biobank, and randomly split the sample into ⅔ training and ⅓ replication splits. For the training samples we perform discovery of vertex-wise measures of area, thickness and sulcal depth as in (4). Having discovered genomic loci in training folds, we perform replication of these loci in the test sets. To perform replication for each SNP we calculate a PolyVertex Score (PVS) (similar to (6,7)) from imaging data in the test set for each MOSTest discovered locus. This PVS aggregates the distributed effects across the cortex by taking a weighted sum across all vertices using mass univariate z statistics as weights from the training set. This approach is similar to the widely used method of Polygenic Risk Scores (PRS) in genetics(8), where instead of predicting a phenotype we are predicting a single genomic variant and instead of using distributed effects across the genome as predictors we use the distributed effects across the cortex, estimated in the training set. For each discovered training set we generate a PVS for each individual, which represents a continuous prediction of the genotype in the test set. We then correlate each PVS with its corresponding measured genomic variant in the test set to test how well these discovered loci replicate (one tailed t test, p<0.05). We test this MOSTest discovery and PVS replication, against an established GWAS approach (min-P)(9). Figure 1 displays a schematic of how replication of how min-P and MOSTest differs for a single discovered variant. We confirm a higher replication yield and comparable replication rate MOSTest versus min-P. Finally, we test the generalization of loci discovered in UK Biobank to a developmental cohort of 9-10 year old children from the Adolescent Brain Cognitive Development^®^ (ABCD; https://abcdstudy.org) Study, where we see a higher yield of replicated loci for MOSTest versus min-P.

**Figure 1.**
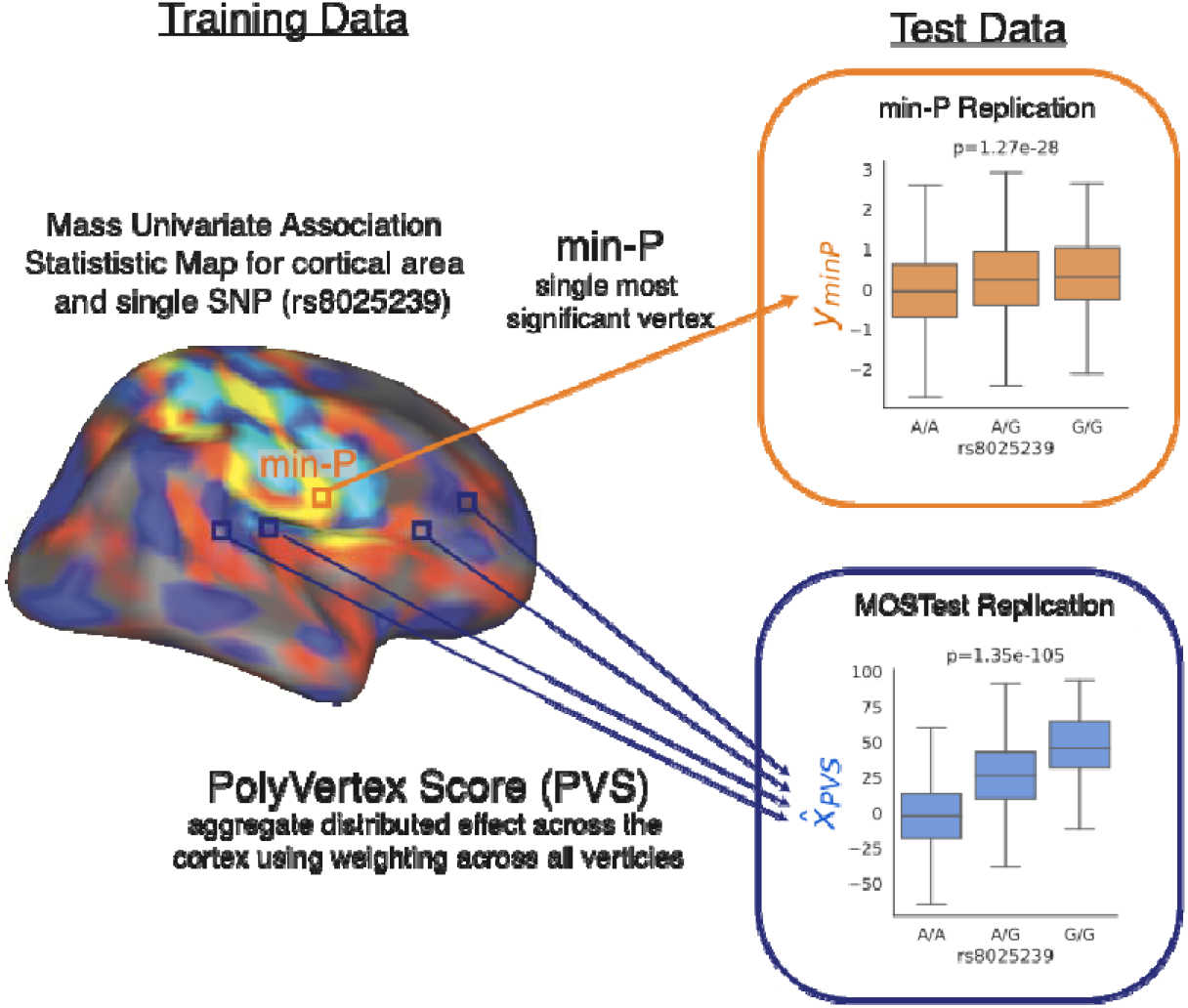
Schematic of replication process for a single SNP. Variant rs8025239 is discovered in training fold and has mass univariate map of association statistics with cortical area. Min-P replication (indicated by orange box and arrow) takes most significant vertex and associates that vertex with variant rs8025239 in test data. MOSTest replication (indicated by blue box and arrow) computes a PolyVertex Score (PVS) in test data which aggregates all effects across the cortex by taking a weighted sum (using association statistics from training set) across all vertices – the PVS is then correlated with the variant rs8025239. This process is repeated for all discovered variants in training set with a separate PVS being generated for each MOSTest discovery. Replication of a variant is defined as p<0.05 in one tailed t-test.

## Results

Across training folds, the UK Biobank sample, we confirm that MOSTest confers up to a 10-fold increase in discovered loci over min-P (area: min-P_yield_=52, MOSTest_yield_=433, thickness: min-P_yield_=42, MOSTest_yield_=367 and sulcal depth min-P_yield_=85, MOSTest_yield_=890). When replication of loci is defined at the nominal level (p<0.05, see methods) we see a higher number of replicated loci, as well as comparable replication rate for MOSTest (area:94%, thickness: 95%, sulcal depth: 95%) vs min-P (area:65%, thickness: 72%, sulcal depth: 80%) – see Figure 2. Averaged across cross-validation folds, we found that the lead SNP of the top locus accounted for more variance in the replication set with MOSTest ( = area:0.037 =3.8×10^−3^, thickness: 0.059 =1.4×10^−2^, sulcal depth: 0.052 =4.0×10^−3^) compared to min-P ( = area: 0.011 =1.2×10^−3^, thickness: 0.011 =2.2×10^−3^, sulcal depth: 0.015 =1.6×10^−3^). If replication is defined more conservatively with significance corrected for the number of discovered loci (p<0.05/# of discovered loci), we again find that MOSTest confers a comparable replication rate (area: 69%, thickness: 70%, sulcal depth: 68%) to min-P (area: 41%, thickness: 55%, sulcal depth: 50%).

**Figure 2.**
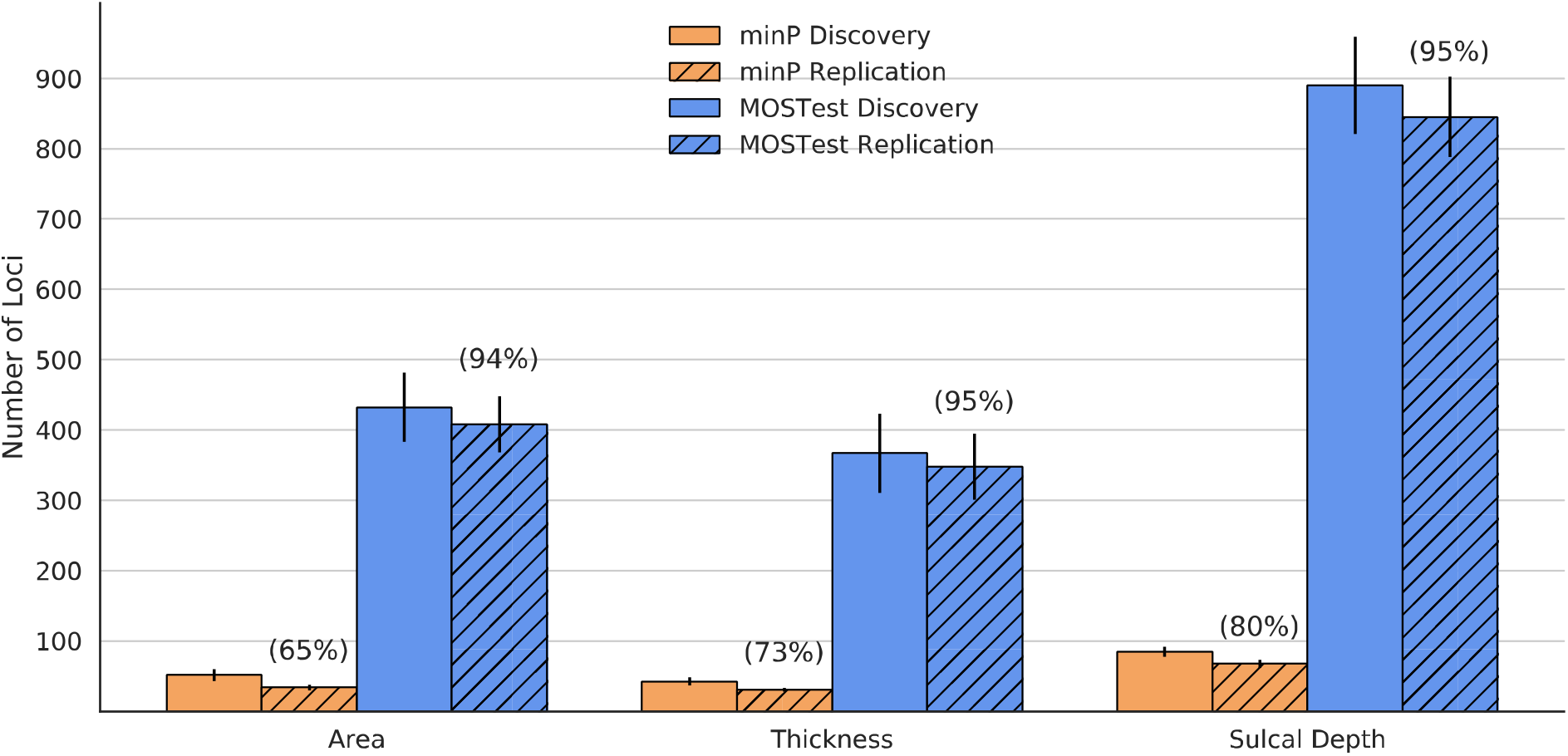
Cross-validation discovery and replication yield within 10-times cross validation within UK Biobank for cortical morphometry measures. Solid bars represent the number of genome wide significant loci associated with each measure. Hashed bars represent the number of loci that replicate in test folds at a nominal significance level (p<0.05). Error bars are standard deviations across 10 cross-validation repetitions. Numbers in parentheses represent replication rate (# of discovered loci / # replicated loci) for each method-phenotype pair.

**Figure 3.**
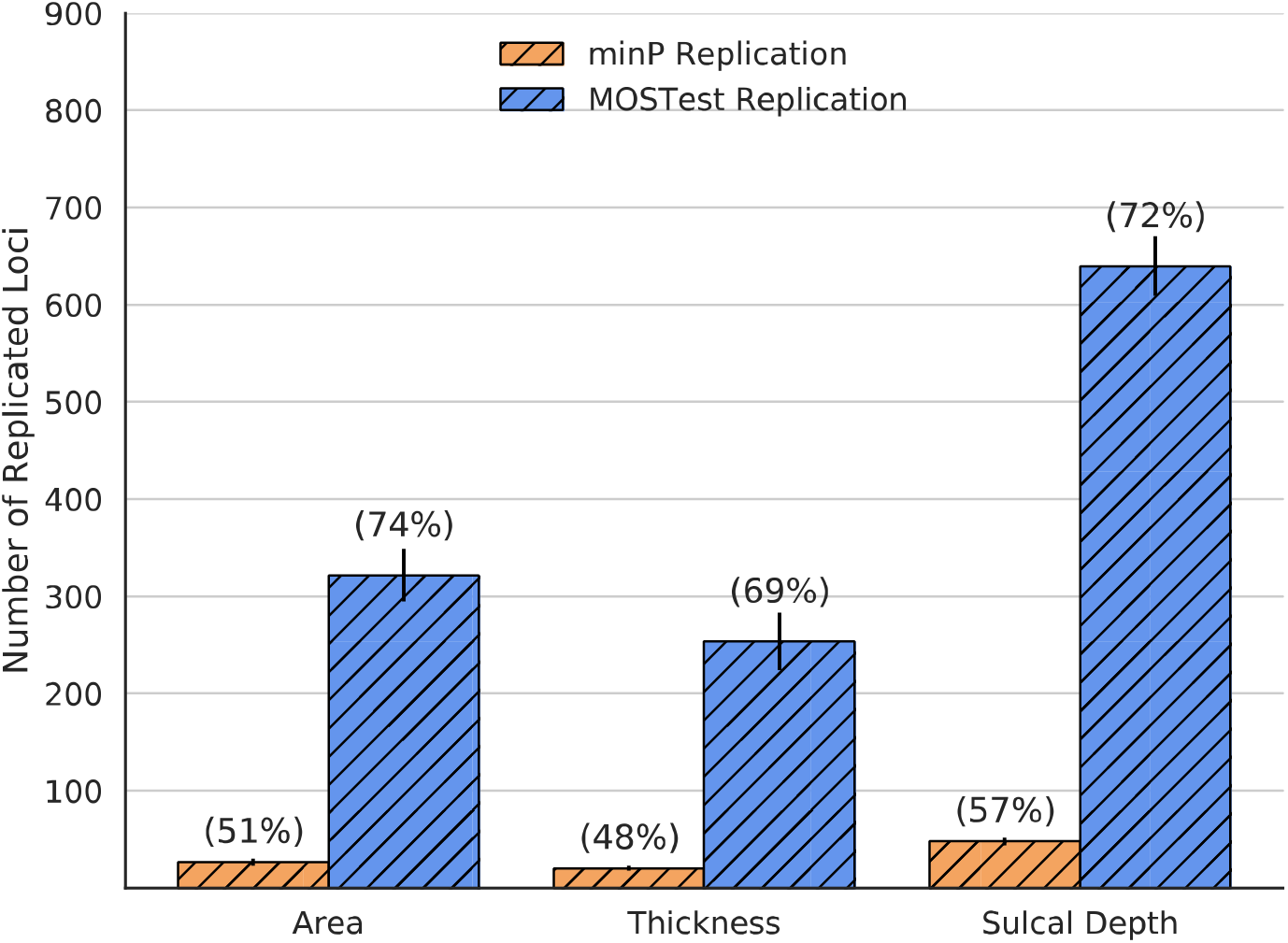
Replication yield within the ABCD dataset across 10 training folds of UK Biobank for cortical morphometry measures. Bars represent the number of loci that replicate in ABCD at a nominal significance level (p<0.05). Error bars are standard deviations across 10 training sets of UK Biobank. Numbers in parentheses represent replication rate (# of discovered loci / # replicated loci). ABCD: Adolescent Brain Cognitive Development Study.

Next, we tested the generalization performance of loci discovered in each training fold of UK Biobank to a developmental cohort of adolescents from the Adolescent Brain Cognitive Development study. Here we once again see a higher absolute number of replicated loci (nominal p<0.05 level), as well as a comparable replication rate for MOSTest (area: 74%, thickness: 69%, sulcal depth: 72%) to min-P (area: 51%, thickness: 48%, sulcal depth: 57%) - see figure 2. Again, the variance explained by the lead SNP of the top locus (averaged across cross-validation folds) accounted for more variance for MOSTest (*R*^2^= area: 1.6×10^−2^ σ=9.0×10^−4^, thickness: 2.9×10^−2^ σ=8.4×10^−3^, sulcal depth: 2.1×10^−2^ σ=1.5×10^−3^) than for min-P (R^2^= area: 7.1×10^−3^ *σ*=1.0×10^−4^, thickness: 4.8×10^−3^ σ=1.1×10^−3^, sulcal depth: 9.1×10^−3^ σ=5.0×10^−4^).

## Discussion

Here we have confirmed the increased power of using a MOSTest across training folds of UK Biobank. Further, through the deployment of PVS we show that loci discovered with MOSTest result in a higher replication yield and comparable replication rate to independent data than established approaches. The comparable replication rate for MOSTest loci (94-95% vs 65-80% for min-P) indicates that the difference in absolute number of replicated loci for MOSTest vs min-P is not merely a result of MOSTest discovering a higher number of loci. Furthermore, we still see a comparable replication rate when we penalize the replication significance threshold by the number of loci discovered by each method (i.e. p<0.05/ # of discovered loci). This underscores the distributed effects of the genome across the cortex, which multivariate methods are better powered to capture and in turn, will display stronger generalization to independent data.

Additionally, we have shown that genetic-cortical morphology associations learned within an adult population (mean age 64 years) of individuals from the UK generalize out of sample to adolescents aged 9-10 years old in the United States of America taken from the ABCD study. There are marked differences between the training sample of UK Biobank and validation sample of ABCD including: large age differences, a high degree of ancestry admixture in ABCD, different scanners used, imaging protocols and the number of individuals in validation sets. In spite of these differences we observe a high replication rate in ABCD of discoveries found within UK Biobank via MOSTest. We see higher replication for cortical area and sulcal depth in ABCD than for cortical thickness. Cortical thickness changes more dynamically over the lifespan(10), therefore, given the large age disparity between the two samples, perhaps it is not a surprise to see that cortical thickness is the measure that exhibits the largest reduction in replication rates in ABCD when compared across cross-validation folds of UK Biobank for MOSTest (69% vs 95%). We may expect that the replication rate of discovered cortical thickness loci to increase as the children develop, a hypothesis that can be tested as more longitudinal ABCD data is collected. Despite differences across these datasets we observe greater replication of UK Biobank discovered loci in ABCD when taking into account the multivariate nature of associations across the cortex (i.e. MOSTest and PVS).

Furthermore, we demonstrated that lead MOSTest discoveries explained a notable amount of variance out of sample, by GWAS standards: 3-6% in UK Biobank and 1-3% in ABCD. Methods, such as MOSTest and PVS, that result in high replication yield and out of sample variance explained may support precision medicine efforts(11). In particular if these methods are deployed on disorders of the brain they may provide complimentary predictive power to well established models such as Polygenic Risk Scores.

The training data used here to detect loci and train PVS projections weights were taken from individuals of European ancestry from the UK Biobank. We may expect that the genetic architecture of cortical morphology to differ between ancestry groups(12). We also acknowledge that our use of PVS to predict genotypes out of sample is just one possible projection weighting scheme, which may not provide optimal out of sample prediction. Here we have demonstrated the high generalization performance of cortical morphology discoveries using MOSTest to independent data. This was shown both within study (UK Biobank) and across studies (UK Biobank to ABCD) despite substantial age differences of participants. This work underscores the importance of deploying well powered multivariate methods when performing GWAS on high dimensional phenotypes, both for discovery and replication.

## Methods

The UK Biobank sample and methods used for min-P and MOSTest discovery overlap with previous work(4,5).

### UK Biobank Sample

Genotypes, MRI scans, demographic and clinical data were obtained from the UK Biobank under accession number 27412, excluding participants who withdrew their consent. For this study we selected white British individuals (as derived from both self-declared ethnicity and principal component analysis) who had undergone the neuroimaging protocol. The resulting sample contained 35,644 individuals with a mean age of 64.4 years (standard deviation 7.5 years), 18,433 female. T_1_-weighted MRI scans were collected from three scanning sites throughout the United Kingdom, all on identically configured Siemens Skyra 3T scanners, with 32-channel receive head coils. We used UK Biobank v3 imputed genotype data(13).

### Adolescent Brain Cognitive Development^®^ (ABCD) Sample

The ABCD study is a longitudinal study across 21 data acquisition site following 11,878 children starting at 9 and 10 years old. This paper analyzed the full baseline sample from data release 3.0 (NDA DOI:10.151.54/1519007). The ABCD study used school-based recruitment strategies to create a population-based, demographically diverse sample with heterogeneous ancestry. T_1_-weighted MRI scans were collected using Siemens Prisma, GE 750 and Phillips 3T scanners. Scanning protocols were harmonized across 21 acquisition sites. Genetic ancestry factors were estimated using fastStructure(14) with four ancestry groups. Genotype data was imputed at the Michigan Imputation Server(15), using the HRC reference panel as described in(16,17). We selected individuals who had passed neuroimaging and genetic quality control checks, resulting in 8,336 individuals with a mean age of 9.9 years (standard deviation 0.62 years), 3,974 female.

### Data processing

T1-weighted structural MRI scans were processed with the FreeSurfer v5.3 standard “recon-all” processing pipeline(18) to generate 1,284 non-smoothed vertex-wise measures (ico3 downsampling with the medial wall removed) summarizing cortical surface area, thickness and sulcal depth. All measures were pre-residualized for age, sex, scanner site, the first ten genetic principal components. In contrast to previous MOSTest work(3,5) we did not pre-residualize for global measures specific to each set of variables (total cortical surface area or mean cortical thickness) as there is no clear corollary for sulcal depth, nor did we control for Euler number. Subsequently, a rank-based inverse normal transformation was applied to the residualized measures. For genomic data we carried out standard quality-checks as described previously(3), setting a minor allele frequency threshold of 0.5% and finding the intersecting variants between UK Biobank and ABCD, leaving 8 million variants. Variants were tested for association with cortical surface area, cortical thickness and sulcal depth at each vertex using the standard univariate GWAS procedure. Resulting univariate p-values and effect sizes were further combined in the MOSTest and min-P analyses to identify area, thickness and sulcal depth associated loci.

### Cross validation

We performed 10 times cross validation within UK Biobank with approximately ⅔ training, ⅓ testing splits, performed randomly except for related individuals were kept together. Due to relatedness in the sample we wanted to ensure that individuals who were highly genetically related were not split across training and testing folds. We estimated relatedness using ‘plink – genome’ and from this defined relatedness clusters with individuals who were related as 3rd degree relatives (threshold>1/8). This resulted in 34,813 clusters. Across the 10 folds, the mean training sample size was 24,471 individuals (S.D. =9.5).

### MOSTest Discovery

Consider *N* variants and *M* (pre-residualized) phenotypes. Let *z_i,j_* be a z-score from the univariate association test between i^th^ variant and j^th^ (residualized) phenotype and ***z**_j_* be the vector of z-scores of the i^th^ variant across phenotypes. Let ***Y*** be a matrix of (pre-residualized) phenotypes with *I* (individuals) rows and *M* (phenotypes) columns, and ***R*** be its correlation matrix. ***R*** can be decomposed using singular valued decomposition as ***R*** = ***USV***^T^ (***U*** and ***V*** – orthogonal matrixes, ***S*** – diagonal matrix with singular values on its diagonal). Consider the regularized version of the correlation matrix ***R*** = ***US_r_V***^*T*^, where ***S***_*r*_ is obtained from ***S*** by keeping *r* largest singular values and replacing the remaining with *r*^*th*^ largest. The MOSTest statistics for i^th^ variant (scalar) is then estimated as 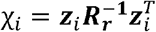, where regularization parameter is selected separately for cortical area, thickness and sulcal depth to maximize the yield of genome-wide significant loci. As established in previous work(3–5) the largest yield for cortical surface area is obtained with *r* =10; the optimal choice for cortical thickness and sulcal depth was *r* =20. The distribution of the test statistics under null 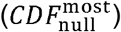 is approximated from the observed distribution of the test statistics with permuted genotypes, using the empirical distribution in the 99.99 percentile and Gamma distribution in the upper tail, where shape and scale parameters of Gamma distribution are fit to the observed data. The p-value of the MOSTest test statistic for the i^th^ variant is then obtained as 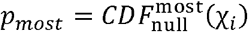.

### min-P Discovery

Similar to the MOSTest analysis, consider *N* variants *M* and pre-residualized phenotypes. Let *z_i,j_* be a z-score from the univariate association test between i^th^ variant and j^th^ (residualized) phenotype and ***z***_*i*_ be the vector of z-scores of the i^th^ variant across phenotypes. The min-P statistics for the i^th^ variant is then estimated as *y*_*i*_ = 2Φ(-*max*_*j=1…M*_(|*z_i,j_*|)), where Φ is a cumulative distribution function of the standard normal distribution. The distribution of the min-P test statistics under null 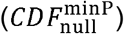 is approximated from the observed distribution of the test statistics with permuted genotypes, using the empirical distribution in the 99.99^th^ percentile and Beta distribution in the upper tail, where shape parameters of Beta distribution (α and β) are fit to the observed data. The p-value of the min-P test statistic for the i^th^ variant is then obtained as 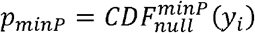.

### Locus definition

Independent significant SNPs and genomic loci were identified in accordance with the PGC locus definition, as also used in FUMA SNP2GENE(19). First, we select a subset of SNPs that pass genome-wide significance threshold 5×10^−8^, and use PLINK to perform a clumping procedure at LD r2=0.6, to identify the list of independent significant SNPs. Second, we clump the list of independent significant SNPs at LD r2=0.1 threshold to identify lead SNPs. Third, we query the reference panel for all candidate SNPs in LD r2 of 0.1 or higher with any lead SNPs. Further, for each lead SNP, it’s corresponding genomic loci is defined as a contiguous region of the lead SNPs’ chromosome, containing all candidate SNPs in r2=0.1 or higher LD with the lead SNP. Finally, adjacent genomic loci are merged if they are separated by less than 250 KB. Allele LD correlations are computed from EUR population of the 1000 genomes Phase 3 data. Obtained clumps of variants were considered as independent genome-wide significant genetic loci.

### Replication of Discovered Variants

A schematic displaying the difference between min-P and MOSTest replication is displayed in Figure 1. For genome-wide significant loci defined in the training folds, we performed replication in test folds of UK Biobank, as well as the whole sample of ABCD. Let ***X***^test^ represent the genotype matrix of individuals in the test set of *I* individuals and *N* variants and ***Y***^test^ represent the phenotype matrix of *I* individuals and *M* (pre-residualized) phenotypes. Replication was performed in one of two ways, depending on whether the genetic variant was discovered using min-P or MOSTest. Firstly, for a min-P discovery, implicated by the association statistic *z_i,j_*, the i^th^ variant, 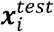, is associated with the j^th^ (residualized) phenotype 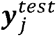, in the test set. Secondly, for MOSTest validation the i^th^ discovered loci corresponds to a vector of mass univariate association statistics across all vertices ***z***_***i***_ - these are used to generate projection weights to create a PolyVertex Score (PVS) (7), 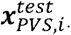. This approach largely mirrors the use of polygenic scores used in genetics, where here we are aggregating effects of vertices across the cortex. For polygenic scores, it is well known that the correlation structure (i.e. linkage disequilibium) across the genome can result in suboptimal out of sample performance. This has motivated techniques like LD-Pred(20) and PRSice(21) to first account for this genomic correlation before generating scores. Similarly, we decorrelate the association statistics, ***z***_*i*_, as 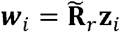 using the regularized correlation matrix 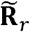 that was learned in the training fold. We then generate the polyvertex score for the i^th^ genomic variant as the dot product of ***w***_*i*_ with the (pre-residualized) phenotype matrix, ***Y***^test^, in the test set: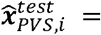 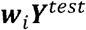.

To perform the association in test set, both of min-P and MOSTest/PVS replications, we used linear mixed-effect models (LMMs) to control for genetic/family relatedness – this is particularly relevant for the ABCD dataset which has a high degree of family relatedness. We used a single fixed effect of the discovered variant, *x*_*i*_, and a random effect intercept using a grouping, *c*, of either: i) genetic relatedness cluster (defined above) for UK Biobank replication or ii) family id (rel_family_id) for ABCD replication. The response variable was either a) the most significant vertex for min-P validation, *y*_*minP, i*_, or b) the computed PVS, 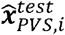, for MOSTest. For min-P replication:

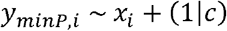

And for MOSTest replication:

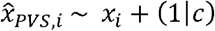

As the phenotype matrix, ***Y***^test^, was pre-residualized for covariates before taking the most significant vertex (min-P) or computing the PVS (MOSTest) we did not need to control for other covariates. We fit an LMM for each discovered locus in training set. For both min-P and MOSTest validation, we one-tailed p values from t statistics of the fixed effect as we assume the effect to be in the same direction for training folds and test sets. To define replicated loci we use a nominal p value threshold of 0.05 for associations. Due to the higher number of discovered loci for MOSTest vs min-P, we additionally report the number of loci validated at a Bonferroni corrected threshold, where this number of independent tests is taken to be the number of discovered loci in the training set. This corrected threshold penalizes MOSTest to a greater extent than min-P for discovering a larger number of loci. We calculate the variance explained by the single lead i^th^ variant in the replication sample from t statistics of *x*_*i*_ from fitted LMMs and degrees of freedom (*df*) as: 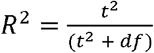 We report the average and standard deviation (σ) of this value across training folds.

## ABCD Acknowledgement

Data used in the preparation of this article were obtained from the **Adolescent Brain Cognitive Development℠ Study** (**ABCD Study®**) (https://abcdstudy.org), held in the NIMH Data Archive (NDA). This is a multisite, longitudinal study designed to recruit more than 10,000 children age 9-10 and follow them over 10 years into early adulthood. The ABCD Study is supported by the National Institutes of Health and additional federal partners under award numbers: U01DA041022, U01DA041028, U01DA041048, U01DA041089, U01DA041106, U01DA041117, U01DA041120, U01DA041134, U01DA041148, U01DA041156, U01DA041174, U24DA041123, and U24DA041147

A full list of supporters is available at https://abcdstudy.org/federal-partners/. A listing of participating sites and a complete listing of the study investigators can be found at https://abcdstudy.org/principal-investigators.html. ABCD Study consortium investigators designed and implemented the study and/or provided data but did not necessarily participate in analysis or writing of this report. This manuscript reflects the views of the authors and may not reflect the opinions or views of the NIH or ABCD Study consortium investigators.

The ABCD data repository grows and changes over time. The ABCD data used in this came from [NIMH Data Archive Digital Object Identifier (10.151.54/1519007)].

